# Reproducible and shareable bioinformatics pipelines from natural-language prompts

**DOI:** 10.64898/2026.05.28.719125

**Authors:** Hyeon-Min Kim, Hwayeon Jeong, Abyot Melkamu Mekonnen, Yeongjun Kim, Youngchul Oh, Heetak Lee, Cheulhee Jung, Jeongbin Park

**Author notes:** To whom correspondence should be addressed: J.P.

## Abstract

Large language models (LLMs) are increasingly used to generate bioinformatics pipelines and to carry out analyses from natural-language prompts. However, the resulting analyses are often difficult to reproduce across sessions, owing to the non-deterministic nature of LLM-driven conversations and heterogeneity of local execution environments, and cannot run on remote high-performance computing (HPC) servers or be shared and reused. We present Autopipe, a platform that guides any Model Context Protocol (MCP) - compatible LLM to produce, execute, and publish source-preserved, re-executable containerized pipelines. Autopipe enables users to execute bioinformatics pipelines on any on-premises remote servers - supported by comprehensive setup documentation aimed at researchers without prior server-administration experience - and to visualize results through an extensible web-based viewer. The Autopipe platform comprises four components: a desktop application with an embedded MCP server for pipeline management and remote execution, an online registry for pipeline and plugin discovery, a web-based result viewer, and a CLI tool for customizing viewer plugins. Autopipe turns conversational analysis into re-executable and shareable workflows. Autopipe is freely available at https://autopipe.org/.

## Main

Modern biology generates massive and heterogeneous datasets across genomic, transcriptomic, and proteomic scales, and their analysis increasingly relies on multi-step computational workflows that combine many specialized tools. To reduce the effort of building such workflows, large language models (LLMs) have been increasingly applied to bioinformatics, both to generate analysis pipelines from natural-language prompts^1, 2^ and to carry out multi-step analyses through conversational, agent-based systems^3–5^. A common way to extend a general-purpose LLM for these tasks is to connect it to domain-specific tools and skill modules, for example, through the Model Context Protocol (MCP) or curated skill collections^6^.

However, current LLM-based approaches face four main limitations for end-to-end bioinformatics analysis. First, the generated analysis code is not inherently preserved as a reusable artifact; once a conversational session ends, the analysis logic is typically lost, and regenerating the same pipeline from an identical prompt may yield a different implementation due to LLM non-determinism. Second, chat-based interfaces cannot perform analysis on on-premises servers, often vulnerable to data privacy concerns. Third, existing AI-based tools have limited support for free access to large binary or tabular scientific data within remote servers, such as BAM, VCF, or H5AD files that are often used in bioinformatics. Fourth, existing AI-driven systems are typically tied to a specific LLM provider, preventing users from leveraging the affordable commercial AI subscriptions they already maintain or selecting the most suitable LLM for their workflow. A range of tools already address individual stages of the bioinformatics workflow lifecycle: workflow management systems support reproducible and scalable execution^7, 8^, the sharing and reuse of published workflows^9, 10^, and the discovery and construction of new ones^11, 12^, providing much of the reproducibility and shareability that LLM-driven analysis lacks. These tools, however, require manual coding and expertise and operate separately from conversational, LLM-driven generation. To overcome these limitations, an LLM’s generation can be guided by a “harness”, in which model outputs are constrained through structured templates and execution scaffolds rather than relying on free-form generation alone. This approach, recently formalized as harness engineering in LLM-based software engineering^13–16^, makes model-driven generation more reliable and reproducible. Guiding an LLM to emit standardized, containerized, and shareable pipelines could thus combine the accessibility of conversational generation with the reproducible execution and sharing that workflow tools provide.

To our knowledge, there is no existing easy-to-use AI-assisted system that extends this harness concept to support automatic reproducible workflow management, execution of workflows, and interactive visualization of large data within an on-premises server environment. Running AI coding assistants directly on such a server system still requires command-line proficiency and the ability to configure execution environments, which remains a significant barrier for researchers lacking any computational background.

To this end, here we present Autopipe, an open platform for bioinformatics workflows that provides a natural language-based end-to-end analysis using any LLM to use, produce, and share re-executable containerized pipelines. Using any existing LLMs that support MCP, one can easily create and manage reproducible pipelines, execute on on-premises servers, and share the workflows via an online registry, by just asking them to AI in their natural language. After running a pipeline, one can easily visualize the analysis results through a web-based viewer, which supports various plugins for visualization of common scientific file formats on servers.

### Autopipe Overview

Autopipe consists of seven steps to generate pipelines from a simple command with natural language: Describe, Generate, Build, Execute, View Results, Upload, and Publish (Fig. 1). To implement this, Autopipe comprises four components: a cross-platform desktop application with an embedded MCP server (Autopipe), a web-based pipeline registry (Autopipe Hub), a result viewer (Autopipe Viewer), and a command-line tool for plugin development and publishing (Autopipe-ext).

**Fig. 1:**
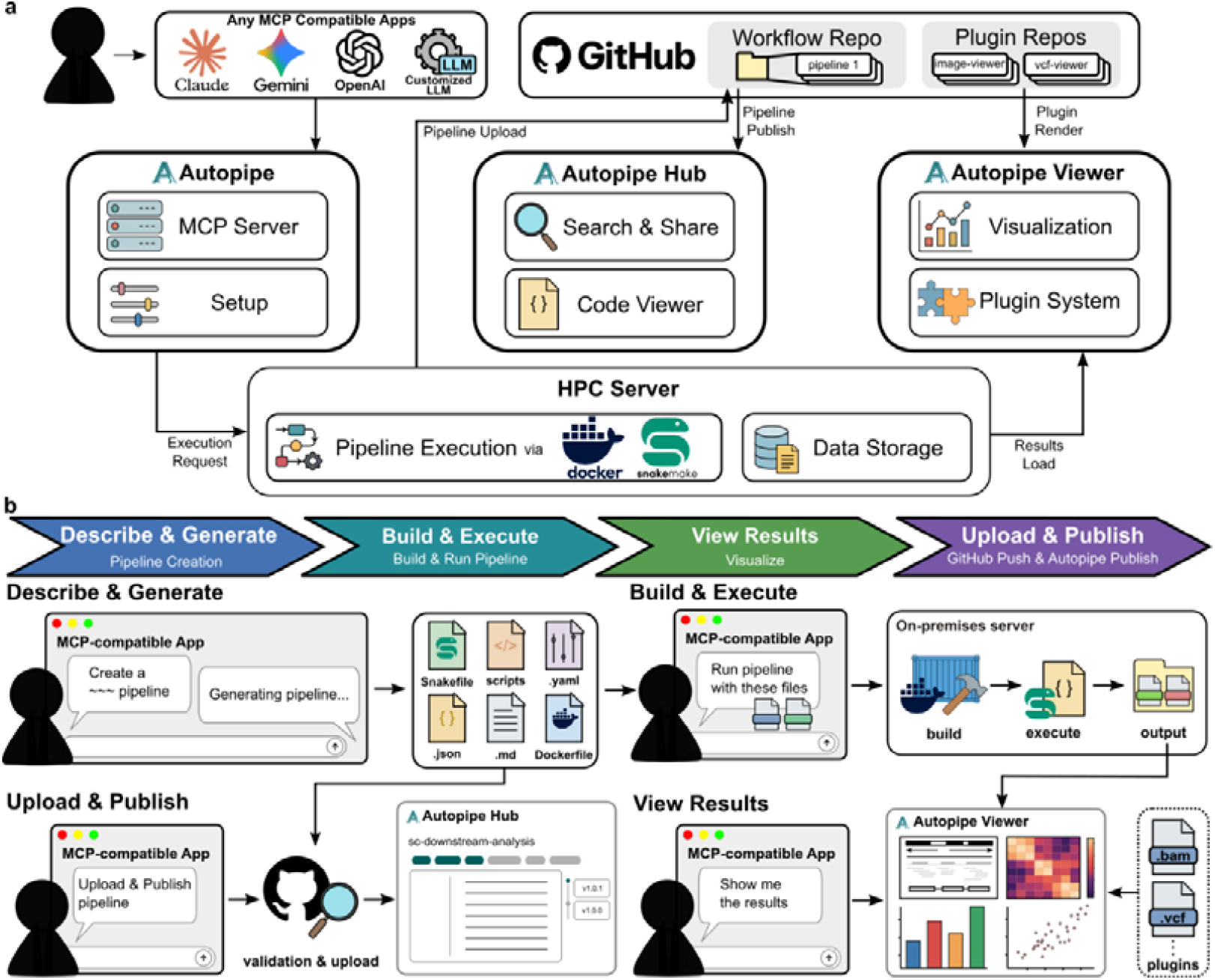
System architecture and end-to-end user workflow of the Autopipe platform. **a**, The Autopipe architecture comprises four core modules. The MCP server and Setup GUI form the central orchestration layer, interfacing with any MCP-compatible AI application (Claude, Gemini, OpenAI, or custom LLMs) to receive natural language requests and invoke the corresponding pipeline management tools. Autopipe Hub handles pipeline search and sharing, and provides a code viewer. Autopipe Viewer renders results via an extensible plugin system, with plugins downloaded from GitHub Plugin Repos and installed locally. Pipeline files are stored in GitHub Workflow Repos and executed on the user’s HPC server via Docker and Snakemake. Arrows indicate data flow between components. **b**, All user interactions are conducted through natural language within an MCP-compatible application. In the Describe & Generate stage, the user describes the desired analysis, and the AI generates a complete pipeline file set per the Autopipe MCP template, including a ‘Snakefile’, analysis scripts, a configuration file, RO-Crate metadata, documentation, and a ‘Dockerfile’. In the Build & Execute stage, Autopipe transfers files to the on-premises server via SSH, builds a Docker image, and executes the pipeline using Snakemake. In the View Results stage, Autopipe Viewer renders outputs in the local browser using format-specific plugins supporting file types such as BAM and VCF. In the Upload & Publish stage, Autopipe commits the pipeline to a GitHub repository and submits it to the public registry, Autopipe Hub.

The MCP server in Autopipe interfaces with MCP-compatible applications such as Claude, Gemini, and Codex, converting analysis requirements described in natural language into containerized pipelines. Since LLMs are reported to prefer widely used programming languages^17^, and this preference is expected to hold for smaller, locally hosted models as well, we selected Snakemake as the pipeline definition language for Autopipe, as it is built on Python.

When a user describes an analysis goal in natural language to an MCP-compatible AI application such as Claude, Codex, or Gemini App, AI uses the tools defined in the Autopipe MCP server to generate a Snakemake pipeline to achieve the given goal, which includes the definition of the workflow structure, tool parameters, and input/output dependencies. The defined workflow is then encapsulated within a Docker^18^ container, with the metadata described in the RO-Crate standard^19^. Users can iteratively refine the generated pipeline through additional natural language instructions until the desired configuration is achieved.

The containerized pipeline is executed within an on-premises server using Docker, and the analysis results are stored in the output directory on the server. Here, we assumed that most end users have a server preconfigured by a hardware service provider; however, for those for whom this is not the case, we provide a shell script that automates the installation of all required dependencies with a single command after login, accompanied by step-by-step documentation for first-time users. After the configuration, the script prints out all the parameters needed for the configuration of the Autopipe desktop app. Upon the execution of the pipeline, the user can request to browse the pipeline outputs. Then, Autopipe Viewer is loaded for visualization of analysis results in the server’s output directory using plugins for each file format.

When the user requests publishing the pipeline, Autopipe uploads the pipeline to a GitHub Repository, using the user’s GitHub credentials. After that, the GitHub repository is registered at Autopipe Hub.

### Autopipe Desktop Application

To begin using Autopipe, users launch the desktop application and configure SSH and GitHub authentication credentials in the provided configuration panels. Upon pressing the save and register button, the Autopipe MCP server is initialized and becomes ready to be connected to MCP-compatible AI applications. Once connected, users can perform all pipeline-related operations – including pipeline generation, building, execution, and result retrieval – through natural language conversations within the AI application, leveraging the tools provided by the Autopipe MCP server (see Supplementary Information). The desktop application also includes a plugin management button, where users can install and update viewer plugins for data-type-specific result visualization.

### Autopipe Viewer and Plugins

The result viewer visualizes analysis outputs directly in the user’s browser without requiring manual file transfer from remote servers (Fig. 2). Thirteen default plugins covering major bioinformatics file types are preinstalled, each dedicated to visualizing scientific data file formats. The default plugins include the formats frequently used in bioinformatics, such as VCF, BED, GFF, BAM, BCF, HDF5, images, PDF, and tabular data, serving examples for further development of new plugins supporting other file types. When a user opens a file, the viewer matches the file extension against installed plugin manifests and delegates rendering to the corresponding plugin. The source code of the plugins is available, which showcases interactive browsing and streaming of large-scale output files, such as multi-gigabyte VCF, BED, and GFF files, directly in the browser. To further assist custom plugin development, we provide Autopipe-ext (see Methods), which provides a boilerplate for a plugin.

**Fig. 2:**
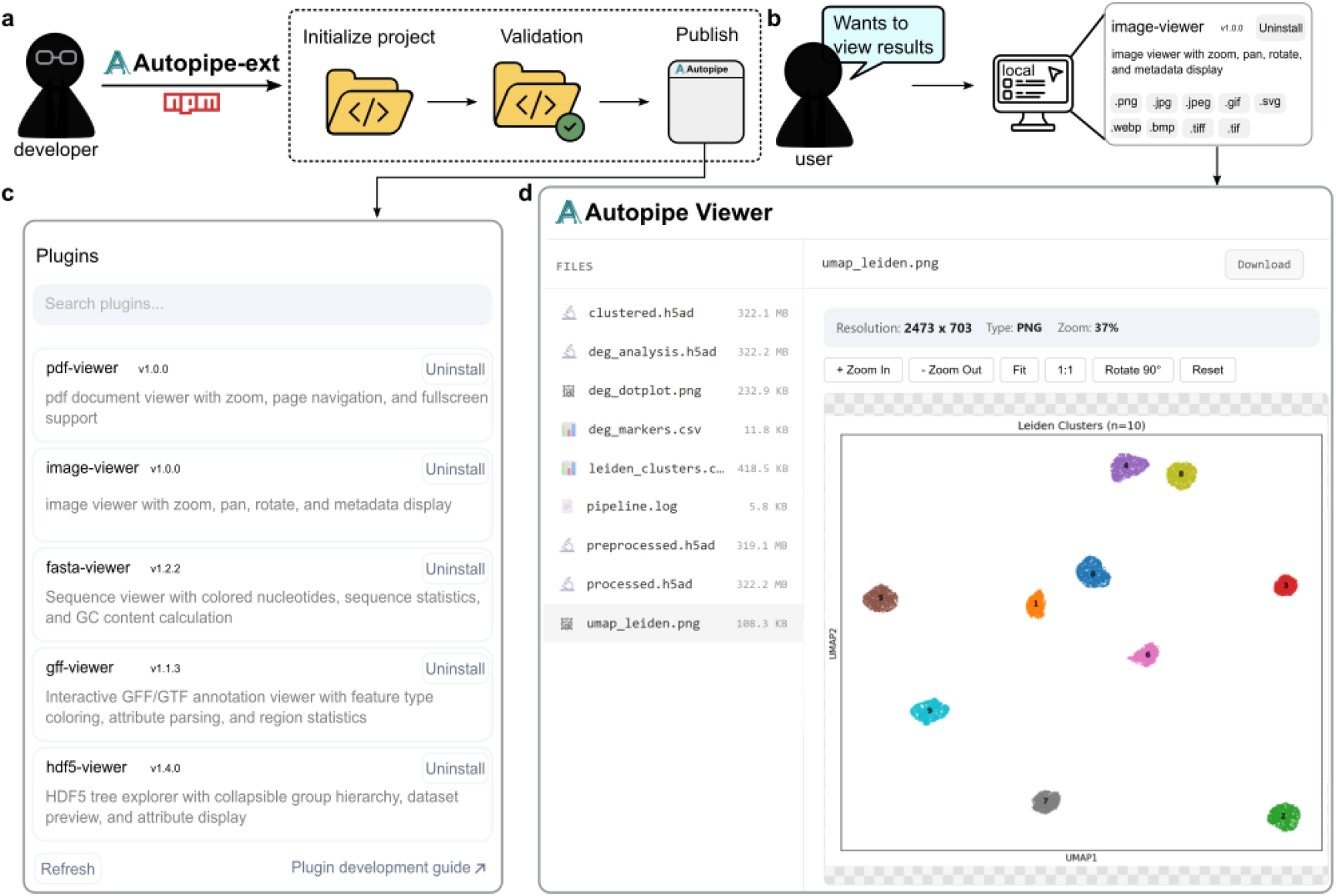
Overview of the Autopipe plugin system and integrated viewer interface. **a**, Developer workflow for creating viewer plugins using the Autopipe-ext CLI. A developer initializes a plugin project, implements the visualization logic, validates the plugin structure, and publishes it to Autopipe Hub, where it becomes available in the Plugins button of the Autopipe desktop application (**c**). **b**, User-side workflow for visualizing analysis results. When a user requests to visualize results, Autopipe Viewer renders the output files in the remote servers in the local browser using installed plugins (**d**). **c**, Plugins panel in the Autopipe desktop application, showing installed plugins handling scientific file formats. **d**, Autopipe Viewer overview. The left panel lists output files. The right panel shows a scatter plot.

### Autopipe Hub

AutoPipe Hub is a web-based registry for pipelines. It stores metadata and GitHub repository URLs and the version-specific published Git tag of deposited pipelines, which includes its name, description, bioinformatics tools used, input and output formats, tags, version, and author information. The pipeline detail view displays this metadata alongside the source code fetched on demand from the tagged GitHub source repository, presented via a file tree and a syntax-highlighted code viewer, supporting source-level inspection before execution.

Upon successful submission of pipelines, the metadata is stored in the PostgreSQL^20^ database and publicly available via the web interface. The web interface of Autopipe Hub provides keyword search and tag-based filtering to aid users in discovering desirable pipelines by pipeline name, description, or tool dependencies. One can describe the desired analysis for AI to recommend one or directly type the pipeline name or ID in natural language, then it will automatically identify the matched published pipelines in Autopipe Hub and execute them in the local environment. In addition, one can easily modify the downloaded pipeline, for example using different tools or parameters, in natural language if needed (Fig. 3).

**Fig. 3:**
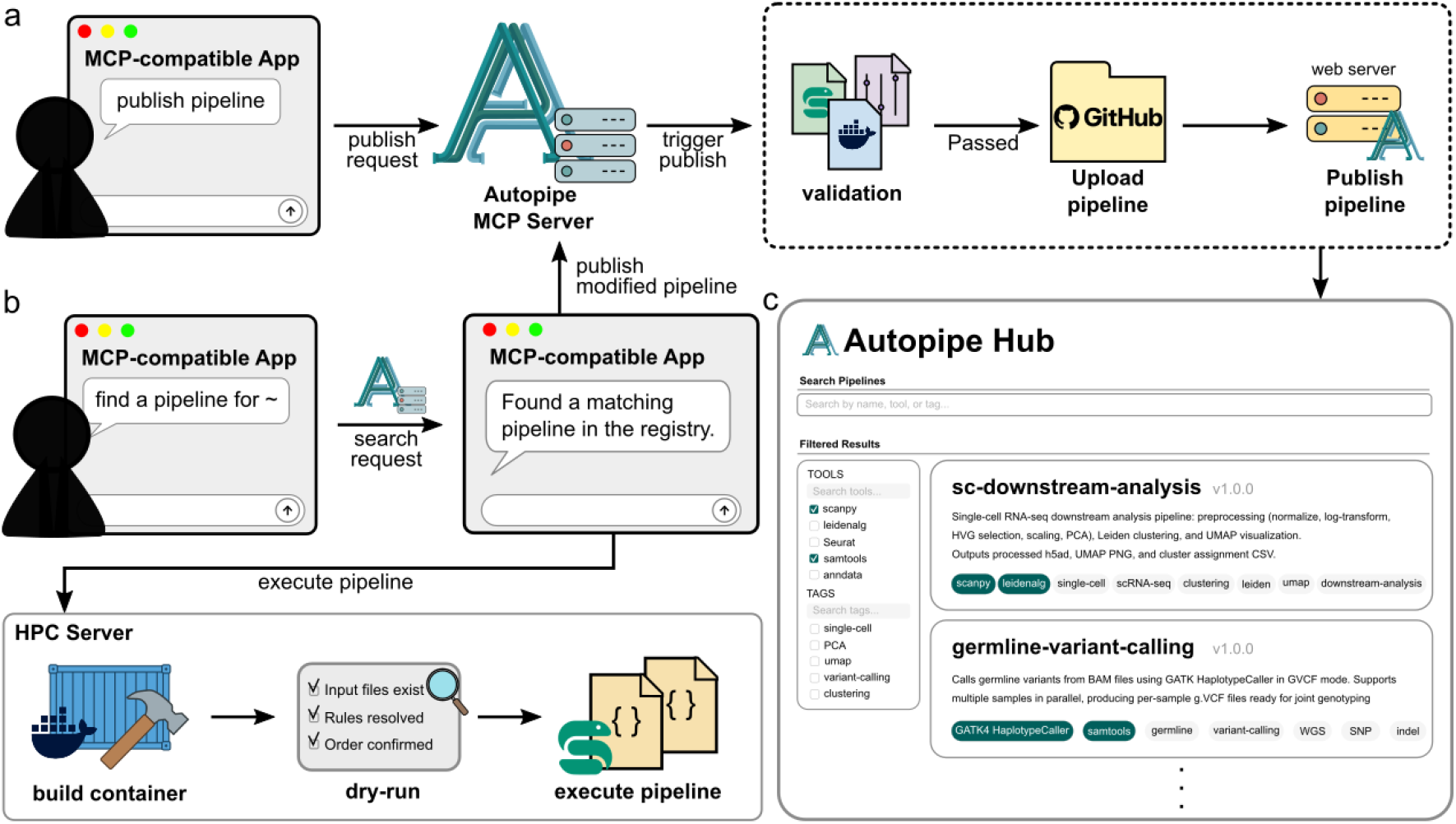
Autopipe Hub pipeline registry workflows and interface. **a**, Pipeline publishing workflow. A user submits a publish request through an MCP-compatible AI app, which triggers the Autopipe MCP server to validate the pipeline files. If validation passes, the files are uploaded to the user’s GitHub repository, and the pipeline metadata is then registered to the Autopipe web server, making the pipeline publicly available on Autopipe Hub. **b**, Pipeline discovery and execution workflow. A user describes a desired analysis through an MCP-compatible AI app, which sends a search request to the Autopipe MCP server. The server queries Autopipe Hub and returns a matching pipeline to the user. The user can modify the retrieved pipeline and publish it back to Autopipe Hub following the workflow described in panel (**a**). Furthermore, the user can execute the pipeline directly, triggering container build on the remote HPC server, followed by a dry-run that verifies input file availability, rule resolution, and execution order. **c**, Autopipe Hub interface. The Autopipe Hub main page displays a searchable and filterable registry of published pipelines. Users can search pipelines by name, tool, or tag, and apply filters using the Tools and Tags panels on the left. Each pipeline entry displays its name, version, description, output file formats, and associated tool and category tags.

**Fig. 4:**
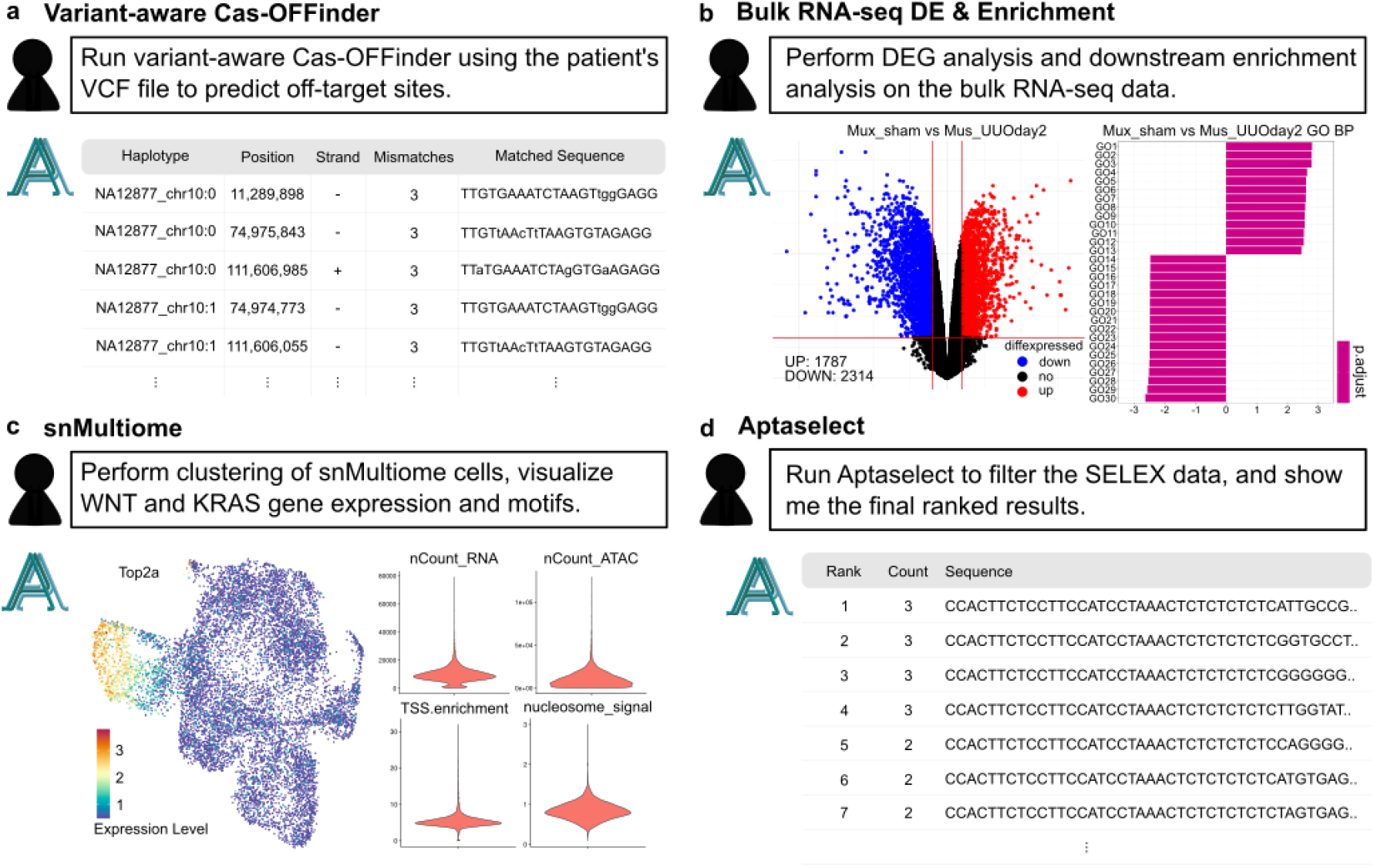
Example pipeline execution results across four bioinformatics scenarios. Each panel shows a short summary of the user’s analysis request (top) with a representative output of the executed pipeline (bottom). **a**, Variant-aware Cas-OFFinder: candidate CRISPR off-target hits called from a phased single-sample VCF, annotated with per-haplotype contig, position, strand, mismatch count, and matched sequence. **b**, Bulk RNA-seq DE and enrichment: a volcano plot of edgeR-based differentially expressed genes (left) and a horizontal bar chart of top GSEA gene sets ranked by normalized enrichment score and colored by adjusted p-value (right). **c**, snMultiome: FeaturePlots of marker gene expression on the integrated UMAP embedding (left) and the pre-filter QC violin plot of the mutant sample showing per-cell distributions of nCount_RNA, nCount_ATAC, TSS enrichment, and nucleosome signal (right). **d**, AptaSelect: paired-end SELEX reads filtered through sequential stages and ranked by frequency to produce the final candidate aptamer list.

### Autopipe can generate, build, execute, and publish bioinformatics pipelines

To evaluate the practical utility of Autopipe, we conducted end-to-end pipeline generation, execution, and publication across four workflows spanning three distinct scenarios: (1) converting a set of existing scripts into a pipeline, (2) reproducing an existing pipeline, and (3) constructing a pipeline for unpublished research. All tests were performed with the Claude Desktop application. The model used was Claude Opus 4.6. Autopipe’s contribution in these examples is the harness rather than the code generation itself, which is performed by the LLM client. The four cases below, therefore, demonstrate the sufficiency of this harness under a current commercial LLM, not a benchmark of LLM-level code-generation correctness.

In the first scenario, we tested whether Autopipe could convert a set of existing analytical scripts into a reproducible pipeline. For this, we asked Autopipe to create a pipeline based on the scripts included in the Variant-aware Cas-OFFinder^21^ tool for CRISPR off-target detection, which is basically a set of scripts and a web frontend, by the tool’s GitHub repository. The pipeline was successfully created, executed, and shared with just four prompts in a single conversational session. Next, we validated the output of the pipeline after execution against that of the original tool using identical input data and parameters. The pipeline produced the same number of off-target sites as the original tool.

In the second scenario, we evaluated Autopipe’s ability to reproduce existing pipelines. We tested 2 different cases for this: (1) a bulk RNA-seq differential expression analysis pipeline, which performs edgeR-based differential expression and enrichment analysis on top of the result of pre-existing nf-core/rnaseq^22^ preprocessing pipeline; and (2) an R-based 10x Genomics single-nucleus multiome (snMultiome) joint analysis^23^ pipeline, which comprises quality control, peak calling, clustering, and visualization using Seurat and Signac. Both pipelines were successfully generated by providing the original analysis scripts and environment configuration files. But in the case of the bulk RNA-seq analysis pipeline, we had to provide the original pipeline’s Dockerfile to the conversational session to resolve the build error; manual code debugging was not needed. Both pipelines produced outputs consistent with their respective original analyses.

In the third scenario, we assessed whether Autopipe could construct a pipeline for entirely novel research. An unpublished pipeline for identifying high-frequency aptamer candidates from SELEX^24^ paired-end sequencing data, named AptaSelect, was generated solely from natural language descriptions of the analytical objectives, without any pre-existing scripts or reference implementations. The pipeline was generated from a prompt containing the algorithm specification, then built and executed through follow-up prompts. The results were inspected through Autopipe Viewer’s CSV-viewer plugin, and the finalized pipeline was uploaded and published to Autopipe Hub. The entire process from generation to publication was completed using 3 prompts in a single conversational session. This case demonstrated that Autopipe can support the development of entirely new analytical workflows.

Several common observations emerged across the four cases. First, errors encountered during build or execution were generally resolved through iterative natural language conversation, with the AI assistant analyzing logs and applying corrections, without requiring manual code debugging. Second, the level of detail in the prompt directly influenced the quality of the generated pipeline. Third, completed pipelines supported subsequent adjustments such as parameter changes and output modifications through conversational interaction, with configuration updates, image rebuilding, and re-execution performed automatically for each adjustment. All four pipelines were registered on Autopipe Hub and are available for search and reuse by other users (see Methods; see Supplementary Information).

### Autopipe supports re-execution of published pipelines

Once a pipeline is generated and published through Autopipe, its source files, metadata, and Dockerfile are associated with a version-specific published Git tag, allowing the published pipeline package to be retrieved independently of the LLM session that produced it. To evaluate the reproducible execution of pipelines generated and published through Autopipe, we retrieved all four pipelines from Autopipe Hub and re-executed them with the same input data used during the original generation. To additionally confirm that the published pipelines re-executed consistently when driven by different MCP-compatible applications, these reproduction examples were performed in the Codex app with GPT-5.5. Each pipeline was identified through natural language requests within the conversational session, and execution was initiated by provisioning input data; Docker image building, dry-run validation, and pipeline execution were performed automatically through conversational interaction (see Supplementary Information). All four pipelines - Cas-OFFinder, bulk RNA-seq differential expression, snMultiome analysis, and AptaSelect – produced the same predefined primary outputs and expected result-file sets, consistent with those obtained during the initial generation runs. The results were verified either through Autopipe Viewer or by direct inspection of the output within the conversational session to confirm primary-output consistency. These reproduction tests indicate that pipelines published to Autopipe Hub support source-preserved, AI-mediated re-execution and reproduce predefined primary results when provided with the same input data and parameters.

### Autopipe enables a local LLM to generate, execute, and publish pipelines

To test whether the Autopipe harness also works with a locally hosted LLM, we connected Autopipe to Qwen 3.6 27B served on an on-premises GPU server through the Open WebUI chat interface (see Methods). Using this local model, a Scanpy-based single-cell RNA-seq downstream pipeline was generated, built, executed, and published to Autopipe Hub entirely through natural-language conversation. The local model required more build-and-execution iterations than the commercial Claude Opus 4.6 client used in the earlier examples, encountering a broader range of container, code, and library-version errors. Nonetheless, every error was diagnosed and corrected by the model through the Autopipe MCP tools, without any human intervention or manual code editing in this case. The workflow ultimately ran to a successful, published outcome. This result suggests that the Autopipe harness can support the same pipeline lifecycle with a locally hosted model; completion of the pipeline lifecycle is carried by the harness rather than by the raw capability of the underlying model.

## Discussion

We presented Autopipe, an open-source, cross-platform desktop application that unifies LLM-assisted pipeline generation, SSH-based on-premises HPC execution, extensible result visualization, and community-driven sharing into a coherent system for bioinformatics workflow management. By embedding a Model Context Protocol server exposing 30 tools for full pipeline lifecycle control (see Supplementary Information), Autopipe enables researchers to describe, generate, build, execute, visualize, and publish Snakemake-based, containerized workflows entirely through natural language interaction.

Our results demonstrate that natural language conversation alone is enough to orchestrate the complete pipeline lifecycle without requiring users to manually navigate separate tools for code generation^1, 2^, workflow execution^7^, registry submission^9^, and data visualization^25^. The combination of Snakemake’s DAG-based execution engine, containerization that encapsulates the execution environment, and RO-Crate 1.1 metadata packaging is designed to make pipelines generated through conversational AI source-preserved and re-executable for identical inputs under tested execution environments, supporting practical reproducibility and FAIR-oriented sharing. The SSH-based remote execution architecture supports data sovereignty for sensitive biomedical datasets by keeping all analyses on user-controlled infrastructure under local credential management and user-approved execution. The JavaScript plugin viewer system and the Autopipe-ext CLI tool empower the community to extend visualization capabilities beyond built-in format support, and the GitHub-backed registry enables execution, visualization, and AI-assisted generation to be shared within a unified ecosystem. Adopting the MCP standard provides interoperability with any MCP-compatible client, establishing a foundation to accommodate the rapidly evolving AI landscape without architectural modifications.

Existing AI-assisted bioinformatics tools^3–5^ and workflow management platforms^8–12, 26^ each address individual aspects of the pipeline lifecycle, such as analysis code generation, workflow sharing, and execution environment provisioning^15^. In the real-world examples presented in this study, this entire process was performed through natural language conversation, effectively removing human effort to manage an on-premises server environment and write source code for pipelines. Combined with the accompanying setup documentation, this design aims to lower the entry barrier for researchers with limited computational background rather than to eliminate it entirely. In our study, pipelines across four different analysis domains were generated, executed, and published entirely via conversation in natural language. Build errors and package dependency conflicts were resolved automatically by the AI assistant, without requiring the user to diagnose or fix the issues. These results suggest that the workflow construction process, which previously required considerable time and expertise across fragmented tools, can be replaced by a single conversational interface.

One concern that may arise is that different LLM clients can produce somewhat different pipeline implementations from the same prompt. Within Autopipe, the harness substantially reduces this variance by constraining outputs to a standardized Snakemake / Docker / RO-Crate structure, and any generated pipeline becomes structured and re-executable because its pipeline logic, configuration, metadata, and container-build specification are preserved in a standardized structure. Beyond this artifact-level guarantee, recent advances in efficient inference and training, such as multi-token prediction and model distillation, are gradually narrowing the performance gap between local and commercial language models. As capable LLMs become viable on user-controlled local hardware, generation itself may become increasingly controllable within a single environment, complementing the workflow-level reproducibility that Autopipe supports through standardized pipelines.

The current limitation of Autopipe is the inability to stream real-time execution progress directly to the LLM client. Because the MCP server operates through a request-response paradigm, users must repeatedly query pipeline status rather than receiving continuous updates during long-running analyses. This polling-based interaction introduces friction, particularly for computationally intensive workflows that may run for hours or days on HPC clusters. However, agentic loop architectures^27^, which allow LLMs to autonomously repeat cycles of executions for next-action determination without user intervention, are becoming increasingly integrated into LLM platforms. Also, agent communication protocols are evolving beyond synchronous request-response toward standards that support asynchronous messaging and streaming^28^. As these architectures and protocols are integrated into LLM, the LLM will be able to autonomously monitor pipeline execution status, detect completion, and notify the user without queries.

Autopipe, as demonstrated throughout this paper, addresses the critical gap between AI-generated analysis and reproducible execution on on-premises remote servers. Autopipe is freely available at https://autopipe.org/.

## Methods

### Autopipe Desktop Application

Autopipe desktop application supports Linux, macOS, and Windows. The desktop application was built using the Tauri framework with a Svelte frontend, providing configuration panels for the SSH connection credentials and the GitHub authentication credentials. These SSH and GitHub credentials are stored locally and never transmitted to the LLM client. For users without a pre-configured remote server, Autopipe provides a shell script that automates the installation of all required dependencies (SSH, Docker, Git, and GitHub CLI) and outputs the exact parameter values needed for the SSH configuration section. The script was validated on a clean Debian environment provisioned as an LXC container with no pre-installed packages, reproducing a fresh server setup via SSH port forwarding and a single command.

The application embeds an MCP server that communicates with LLM clients through either the stdio transport protocol or the Streamable HTTP transport, depending on the client. The MCP standard defines a structured interface through which AI assistants can discover and invoke external tools, enabling bidirectional communication between the LLM and the Autopipe system. When an MCP-compatible client such as Claude Desktop connects to Autopipe, it receives a catalog of available tools along with their input schemas and descriptions, allowing the LLM to select and invoke appropriate tools in response to natural language user requests. To confirm that Autopipe operates well across MCP-compatible applications, we additionally verified that it performs correctly when connected to Gemini CLI and Codex app in place of Claude Desktop.

The MCP server exposes 30 tools organized across 9 functional categories. Workspace tools provide path queries and SSH connection information for the remote server. Pipeline registry tools enable searching, downloading, uploading, validating, publishing, and unpublishing pipelines against Autopipe Hub. Docker build tools handle image building and build status checking on the remote server. Execution tools support pipeline runs, dry-run validation, status monitoring of running jobs, and cleanup of failed executions. Remote file management tools provide reading, writing, listing, downloading, and symbolic link operations on the HPC server. Input preparation tools support downloading files from remote URLs and staging data into the pipeline input directory. Template tools provide ‘Snakefile’ and ‘Dockerfile’ generation guides and file templates for new pipeline creation. Plugin management tools enable listing and browsing locally installed viewer plugins (see Supplementary Information). Visualization tools open analysis results in Autopipe Viewer. The viewer renders analysis results through a local HTTP server with an extensible plugin system (see Methods, “Autopipe Viewer”).

To guide LLMs toward generating well-structured pipelines, the MCP server provides a pipeline template consisting of four files: a ‘Snakefile’ defining the workflow rules and dependencies, a ‘Dockerfile’ specifying the containerized execution environment with all required software dependencies, a ‘config.yaml’ file parameterizing the pipeline inputs and settings, and a ‘ro-crate-metadata.json’ file describing the pipeline metadata in RO-Crate 1.1 format. When a user requests a new pipeline, the template tool returns these base files, and a separate generation guide tool provides the LLM with concrete instructions on how to write the pipeline. This template-driven approach acts as a harness that directs LLM output toward standardized, executable pipeline structures, addressing the reproducibility limitations observed in unconstrained LLM code generation^1, 2, 4^. Following initial configuration through the Autopipe desktop application, users can manage the full pipeline process – from creation through execution, result inspection, and community sharing – through natural language interaction within a single MCP-compatible LLM client alone.

### SSH-Based Remote Execution and Containerized Pipeline Execution

Rather than requiring users to manually log into on-premises servers, Autopipe establishes SSH sessions from the local desktop to remote servers. When a pipeline is submitted, Autopipe transfers the pipeline execution code to the remote server, builds the Docker image, and executes the pipeline within a container. Pipeline executions run as containers on the remote server independently of the local SSH session, ensuring that local network interruptions do not terminate long-running analyses. Input data directories are mounted as read-only volumes to prevent accidental modification of original datasets, consistent with established practices for reproducible containerized workflow execution. Output directories are mounted with write permissions, and all intermediate files and results are written to designated locations. This separation of read-only inputs from writable outputs ensures data integrity throughout the analysis process. In addition to file-system separation, Autopipe follows a user-approved execution model for generated and downloaded pipelines. SSH and GitHub credentials are stored in the local desktop application, and the LLM client invokes MCP tools without directly receiving raw credentials. Before building or executing, the generated or downloaded pipeline remains available for review through the conversational client before any build or execution step is initiated. Generated or registry-downloaded pipelines are treated as executable third-party code: their source files remain visible before build and execution, the LLM client does not receive raw SSH or GitHub credentials, input directories are mounted read-only, and outputs are restricted to designated output directories.

### Autopipe Viewer and Remote Data Streaming

The viewer runs as a local HTTP server on a dynamically assigned port using the Axum framework. Upon installation, 13 default plugins supporting major bioinformatics file formats, including VCF, BED, GFF, BAM, BCF, HDF5, images, PDF, and tabular data are automatically installed from the registry. IGV.js^29^ was used for visualization for genomic data formats (CRAM/BAM/BED/FASTA/GFF).

Result files stored on remote servers are accessed on demand through SSH. For tabular and text-based formats, the viewer executes text stream extraction commands on the remote server via SSH to retrieve only the requested range of rows, returning data in fixed-size pages without transferring entire files. For binary formats such as BAM and CRAM, the viewer relays HTTP range requests from the browser to the remote server through SSH connection, enabling random-access streaming of large files without downloading them in full.

The available rendering formats depend on the plugins installed by the user. When a user opens a file, the viewer matches the file extension against installed plugin manifests and delegates rendering to the corresponding plugin. Community viewer plugins are third-party code; Autopipe validates plugin structure and manifests, but users should install plugins from trusted sources or after source review. Custom plugins can be developed and published using Autopipe-ext (see Methods, “Autopipe-ext CLI Tool”).

### Autopipe-ext CLI Tool

Autopipe provides a Node.js-based command-line interface (CLI) tool, Autopipe-ext. This supports the complete viewer plugin development lifecycle. To create a new plugin, a developer can run the ‘autopipe-ext init’ command. This asks the plugin name, description, and target file extensions in interactive prompts, then generates a boilerplate of the pipeline consisting of the ‘manifest.json’ configuration file, a JavaScript entry file, and a ‘README.md’. The ‘autopipe-ext package’ command validates the plugin directory structure and manifest integrity, verifying that the manifest conforms to the expected schema and the existence of a valid entry file. The ‘autopipe-ext publish’ command automates registry submission by detecting the GitHub remote URL from the local Git configuration, checking whether a plugin with the same name already exists in the registry, and if so, prompting the user to select an appropriate version increment based on the scope of changes. Once published, plugins become available at https://hub.autopipe.org/ and can be installed in the ‘Plugins’ button of the Autopipe desktop application. Overall, the Autopipe-ext CLI Tool allows researchers to develop custom viewers for any file formats specific to their analysis tools or instruments and share them with the community (Fig. 2).

### Publication of the pipeline

The publication pipeline proceeds through three stages. First, users authenticate via GitHub Device Flow OAuth, which happens once at the configuration after the initial installation of the Autopipe desktop app. Second, all files required to execute the pipeline – including Snakefile, Dockerfile, ‘config.yaml’, ‘ro-crate-metadata.json’, ‘README.md’, and any auxiliary scripts or configuration files – are uploaded to a GitHub repository using user-provided credentials. At publication, Autopipe creates and records a version-specific published Git tag for the pipeline, while container images are rebuilt from the deposited Dockerfile during execution. Third, after the user requests publication of the pipeline, Autopipe registers the pipeline metadata in the hub database and links it to the GitHub repository.

### Example 1 - Variant-aware Cas-OFFinder

To validate Autopipe’s end-to-end workflow, we repackaged the analytical workflow of Variant-aware Cas-OFFinder, a CRISPR potential off-target site detection tool that incorporates individual genomic variants from VCF files, into the Autopipe framework.

Pipeline generation was initiated with a prompt referencing the GitHub source repository. The AI assistant read the CLI source code (vcf-cas-offinder_cli.py) to identify the exact tool flags and normalization chain, then automatically generated five pipeline files: a ‘Snakefile’ encoding six rules, a ‘Dockerfile’, ‘config.yaml’, ‘RO-Crate metadata’, and a ‘README.md’. The ‘Snakefile’ faithfully reproduced the original tool’s normalization chain (vcfallelicprimitives | bcftools norm -m- | vcfcreatemulti), per-chromosome VCF splitting to avoid vcf2fasta segfaults, FASTA generation via vcf2fasta -f <ref> -p <prefix> -n NAN, and per-file Cas-OFFinder execution — all matching the original CLI implementation. Input data preparation, including symlink creation for the VCF file and reference genome directory, and handling of plain gzip VCF via zcat | bgzip recompression, was performed automatically.

Docker image building leveraged cached conda layers with pinned versions (vcflib=1.0.3, bcftools=1.21, cas-offinder=2.4.1), and the dry-run validation confirmed a correct DAG of 76 jobs across 24 chromosomes. The first full execution completed all steps without errors but produced empty output because the VCF used UCSC chromosome naming (chr1, chr2, …) while the config listed bare Ensembl-style names (1, 2, …). AutoPipe identified the mismatch by inspecting the split VCF headers and the vcf2fasta error log (“unable to find FASTA index entry for””), corrected the chromosome list in config.yaml, and re-executed. A second iteration further narrowed the chromosome list to only chr6 and chr10 to match the test sample’s actual content, successfully generating four allelic FASTA files (~135–173 MB each, two haplotypes per chromosome).

The initial Cas-OFFinder execution then failed with a PAM/guide length mismatch error (“The length of target sequences should match the length of the pattern sequence”). The original PAM pattern was 17 characters while the guide RNA target was 22 characters; Autopipe identified this from the cas-offinder stderr log and corrected the PAM to NNNNNNNNNNNNNNNNNNNNGG (22 characters), matching the guide length. The subsequent run completed successfully, finding 6 off-target sites across both haplotypes of chromosome 10. Two additional guide RNA sequences (TTGTGAAATCTAAGTGTAGNNN and CTTCACAATTATTCGCCCANNN) were then tested through single-prompt re-executions, with Snakemake automatically reusing the cached normalization and FASTA generation steps and only re-running the cas-offinder and result-combination rules. The final guide produced 8 off-target sites spanning both chr6 and chr10, with position differences between haplotypes confirming that variant-aware coordinate shifts were correctly captured.

After validation, the config was updated to the full chr1–chrY chromosome set for general use, and GitHub upload, and Autopipe Hub publication were requested. The pipeline was uploaded to GitHub and registered in the Autopipe Hub (registry identifier 78). The entire process — from generation through iterative debugging, multi-guide validation, and publication — was completed in a single conversational session. All file generation, dependency resolution, container management, execution, result inspection, and registry publication were performed automatically through Autopipe’s MCP tool interface.

### Example 2 - Bulk RNA-seq Differential expression analysis

To construct a reproducible Bulk RNA-seq analysis workflow, Autopipe was used to generate a pipeline based on a natural language prompt. The pipeline was constructed as a modified version of a previously published Bulk RNA-seq differential expression analysis pipeline.

For data preprocessing, including read quality control, alignment, and quantification, we implemented a pipeline directly following the well-established nf-core/rnaseq workflow (STAR–Salmon), and organized it to allow execution with simple file inputs and a structured description of the workflow. The downstream analysis workflow was designed to perform differential expression and enrichment analysis using edgeR, taking as input preprocessed outputs from the nf-core/rnaseq pipeline (STAR–Salmon workflow)^22^. Specifically, gene-level quantification files (.sf) and a transcript-to-gene mapping file (tx2gene.tsv) were used as inputs. Pipeline generation was initiated by providing the directory containing the original analysis pipeline. This directory included step-by-step functional scripts along with a README file describing the structure and execution of the pipeline. Based on the initial prompt, Autopipe generated the required workflow components, including a ‘Snakefile’, ‘Dockerfile’, and associated scripts.

Environment-related issues arose during the construction of the bulk RNA-seq pipeline. For the preprocessing pipeline, two compatibility issues: a Nextflow version mismatch with the legacy if-block syntax in nf-core/rnaseq v3.22.2’s nextflow.config and the misinterpretation of --genome null as a string literal. Notably, both were diagnosed and resolved by Autopipe entirely on its own. The LLM client identified each failure from the build and runtime logs, applied the corresponding fixes directly to the pipeline files, and re-triggered the build until execution succeeded, all without any manual code editing, debugging, or additional user input. For the downstream DE analysis pipeline, a build failure occurred in the R-based Docker environment due to a dependency issue with the clusterProfiler package. This issue was resolved when the user provided the ‘Dockerfile’ from the original analysis environment, enabling accurate and stable reconstruction of dependencies. This constitutes a manual intervention in which the user supplied an external reference file to guide the AI assistant’s error resolution.

For validation, dry-run testing and full pipeline execution were performed using publicly available datasets^30^. The pipeline produced the same primary outputs as the prior analysis. To evaluate the flexibility of the generated workflow, output modification was performed by leveraging the structured pipeline components produced by Autopipe. An updated R script was generated by Claude AI based on the organized pipeline functions. The script was then executed on the analysis server using Autopipe, and the results were successfully obtained. This process required no additional configuration.

Following validation, the pipeline was successfully uploaded to a personal GitHub repository and published on the Autopipe Hub (registry identifier: 77, 96). The entire process, from pipeline generation to publication, was completed within a single conversational session using only 5 prompts.

### Example 3 - snMultiome

To construct a reproducible single-nucleus multiome (snMultiome) analysis workflow, Autopipe was used to generate a standardized pipeline based on a natural language prompt. The pipeline was built by leveraging pre-existing, manually curated analytical scripts from a published gastric cancer study^23^.

The workflow was designed to perform comprehensive single-cell multi-omic analysis using the Seurat and Signac packages. Specifically, the pipeline was structured to sequentially execute data loading and quality control, peak calling, clustering, differential expression analysis, motif enrichment testing, and downstream visualization.

The pipeline generation was initiated by directly providing the AI assistant with the raw R scripts and environment configuration files as attachments. A detailed prompt was utilized to define the required step-by-step workflow – separating data loading and quality control from visualization steps and instructing the assistant to extract implementation logic, filtering thresholds, and parameters from the attached files. Based on a prompt, Autopipe generated standardized workflow components, including the ‘Dockerfile’, ‘Snakefile’, ‘config.yaml’, ‘README.md’, ‘ro-crate-metadata.json’, and associated R scripts.

Prior to pipeline generation in a Windows environment, manual configuration was required to establish the MCP connection. The JSON file was manually routed to the local Claude Desktop application directory. Furthermore, to resolve directory parsing errors during SSH connection, the default SSH shell on the host machine was modified from PowerShell to WSL Bash via a registry update.

For validation, the generated pipeline was executed, and runtime errors were identified and resolved iteratively through the Autopipe interface. Specifically, deprecation of the aes_string() function in newer ggplot2 versions, the requirement for JoinLayers() in Seurat v5 prior to differential expression testing, and incompatibility of JoinLayers() with ChromatinAssay objects were each identified during execution and corrected through conversational troubleshooting. Snakemake’s caching mechanism allowed re-execution to resume from the failed step, avoiding redundant recomputation of completed upstream steps. The final pipeline produced all expected outputs, including QC reports, UMAP visualizations, differential expression results, and motif enrichment analyses.

Following validation, the finalized pipeline was published to GitHub and the Autopipe Hub (registry identifier #74). All processes, from script aggregation and environment construction to execution, debugging, and registry publication, were facilitated entirely through Autopipe’s conversational interface.

### Example 4 - Aptaselect

AptaSelect is designed to identify enriched aptamer candidate sequences from paired-end FASTQ files generated in SELEX experiments^31, 32^. Unlike previous examples that reconstruct existing workflows, this case evaluates Autopipe’s ability to generate a new analysis pipeline from scratch, solely based on an algorithm specification.

We designed the pipeline dedicated to the structural design of the aptamer library. The library consists of three parts: left/right constant regions, and a central variable region (N-loop).

The analysis proceeds through four major steps: read joining, constant region filtering, candidate extraction, and frequency-based ranking.

In the read joining step, paired-end reads (R1 and R2) are joined into a single sequence based on their overlapping regions. At mismatched positions within overlapping regions, the base with the higher Phred quality score is selected to improve sequence accuracy.

In the constant region filtering step, joined reads are screened to retain only those containing matches to both left and right constant regions, allowing a user-defined number of mismatches to accommodate sequencing errors. After this step, the remaining reads are considered as candidates for a valid aptamer library.

In the candidate extraction step, two sequential extraction modes are applied to the filtered reads. Variable-length extraction first identifies reads containing both constant regions with up to one mismatch allowed in each, and extracts the intervening variable region regardless of its length. Fixed-length extraction is then applied to these results, retaining only reads whose variable region length matches a user-specified value. This stepwise refinement yields two nested candidate sets, a broader variable-length set and a fixed-length subset, allowing users to evaluate enrichment patterns across variable region lengths before focusing on the target length of interest.

In the frequency-based ranking step, the variable region sequences from each extraction mode are independently aggregated and ranked by occurrence frequency. This dual-ranking approach allows users to assess whether high-frequency candidates are robust across both matching conditions or uniquely enriched under mismatch-tolerant conditions, providing a more comprehensive view of the SELEX enrichment landscape. Pipeline generation was initiated using a natural-language prompt describing the algorithm and input data paths. Autopipe automatically generated all required components, including a Python implementation script, ‘Snakefile’, ‘Dockerfile’, a configuration file, RO-Crate metadata, and documentation. The generated pipeline was executed on test data (~10,000 read pairs), and iterative refinement, such as modifying the output format from top-N to full ranking, was achieved through conversational interaction. The final results were inspected through the integrated viewer and successfully published via the Autopipe registry.

The results demonstrate that AptaSelect produces structured and interpretable outputs consistent with SELEX enrichment principles. Together, these results show that Autopipe can translate high-level algorithm descriptions into reproducible pipelines that generate biologically meaningful outputs.

### Pipeline Reproduction Examples

For each pipeline, we assessed re-execution by comparing primary output metrics against the original generation run. To evaluate cross-client re-executability of published pipelines, all four pipelines were re-executed in the Codex desktop app using the GPT-5.5 model and compared with the original generation runs on their primary outputs and result-file sets. Each pipeline was downloaded and re-executed on the on-premises server with the same input data as the original generation runs, and outputs were inspected through Autopipe Viewer or direct file listing within the conversational session. For Variant-aware Cas-OFFinder, the reproduction produced matching off-target site counts, six off-target sites for the first guide setup, and eight off-target sites for the second guide. For the bulk RNA-seq workflow, the preprocessing stage completed all 236 tasks consistent with the original generation run, and the downstream differential expression stage produced the same primary DEG output file. For snMultiome, the reproduction produced an integrated 10,809-cell Seurat object matching the cell count of the original generation run and the same set of result files. For AptaSelect, the reproduction produced identical per-stage unique sequence counts across the four sequential stages on 10,000 read pairs, identical to the original generation run. Reproduction was evaluated at the level of predefined primary outputs and result-file sets, rather than byte-level checksums of all generated files.

### Running Autopipe with a Local LLM

We connected Autopipe to a locally hosted Qwen 3.6 27B model, specifically the Q6_K-quantized unsloth/Qwen3.6-27B-GGUF build, served on an on-premises GPU server. The model was served through llama.cpp’s llama-server with a 262,144-token context window, flash attention, context shifting, and turbo3 key/value cache quantization, exposing an OpenAI-compatible API, and the Jinja chat template was enabled so that the model could emit standardized tool calls. Open WebUI served as the chat interface, with the local model registered as an OpenAI-API connection. The Autopipe MCP server was then registered in Open WebUI’s Manage Tool Servers panel as a streamable HTTP MCP connection, with native function calling enabled.

Pipeline generation was initiated with a natural-language prompt requesting a Scanpy-based single-cell RNA-seq clustering pipeline for a mouse dataset. The model generated a Snakefile, Dockerfile, config.yaml, RO-Crate metadata, a README.md, and Python analysis scripts covering quality control, normalization, highly-variable-gene selection, PCA, Leiden clustering, and a UMAP embedding colored by cluster.

Docker image construction and pipeline execution surfaced a sequence of errors that the model resolved iteratively, working only from the build and runtime logs. These spanned container-build issues, including an unavailable conda package, a NumPy source-build failure, and a shell-symlink error, which the model addressed by installing Snakemake via pip, pinning packages with pre-built wheels, and removing the shell-symlink replacement. They also included a variable-naming conflict in the clustering script, a missing scikit-misc dependency required for Seurat-v3 highly-variable-gene selection, and several Scanpy API changes across versions affecting quality-control metric computation and the PCA and UMAP plotting interfaces. Each fix was applied directly to the pipeline files through the MCP write-file tool, the image was rebuilt, and Snakemake’s caching allowed re-execution to resume from the last completed step, with no manual code editing or human intervention required at any point.

The pipeline was completed on a simulated mouse dataset and produced QC, PCA, and clustering plots, a per-cell QC metrics table, and a text summary report. The model then uploaded the pipeline to GitHub and published it to Autopipe Hub as registry identifier 105, completing the generation-to-publication lifecycle through the local model connected to the Autopipe.

## Declarations

### Ethics approval and consent to participate

Not applicable

### Consent for publication

Not applicable

### Data availability

The Autopipe is freely available at https://autopipe.org/. The source code is available at https://github.com/autopipe-org/autopipe.

### Author contributions

H-MK and JP conceived the study. H-MK developed the Autopipe and designed the figures. HJ, AMM, YK, and YO conducted rigorous testing and provided critical comments. All authors wrote the paper and approved the manuscript.

## Acknowledgements

During the preparation of this manuscript, the authors used the Scientific Paper Writing Assistant (SPWA; https://research.pnucolab.com/spwa)^33^ for structuring and drafting the manuscript, and AI-based language models for reviewing and improving the readability of the text. AI-based coding assistants were also used for software development and debugging. All AI-generated output was reviewed and edited by the authors, who take full responsibility for the content of this publication.

## References

1. Ma, P. et al. Prompt-based bioinformatic pipeline generation for a multi-step metaviral workflow. Bioinformatics Advances 6, vbaf308 (2026).

2. Alam, K. & Roy, B. From Prompt to Pipeline: Large Language Models for Scientific Workflow Development in Bioinformatics. Preprint at 10.48550/arXiv.2507.20122 (2025).

3. Zhang, M. et al. PromptBio: A Multi-Agent AI Platform for Bioinformatics Data Analysis. Preprint at 10.1101/2025.07.05.663295 (2025).

4. Mehandru, N. et al. BioAgents: Bridging the gap in bioinformatics analysis with multi-agent systems. Sci Rep 15, 39036 (2025).

5. Xin, Q. et al. BioInformatics Agent (BIA): Unleashing the Power of Large Language Models to Reshape Bioinformatics Workflow. Preprint at 10.1101/2024.05.22.595240 (2024).

6. K-Dense Inc. Claude scientific skills: a comprehensive collection of scientific tools for Claude AI. GitHub https://github.com/K-Dense-AI/claude-scientific-skills (2026).

7. Di Tommaso, P. et al. Nextflow enables reproducible computational workflows. Nat Biotechnol 35, 316–319 (2017).

8. Galaxy Community. The Galaxy platform for accessible, reproducible, and collaborative data analyses: 2024 update. Nucleic Acids Res. 52, W83–W94 (2024).

9. Gustafsson, O. J. R. et al. WorkflowHub: a registry for computational workflows. Sci Data 12, 837 (2025).

10. Grayson, S., Marinov, D., Katz, D. S. & Milewicz, R. Automatic Reproduction of Workflows in the Snakemake Workflow Catalog and nf-core Registries. In Proceedings of the 2023 ACM Conference on Reproducibility and Replicability 74–84 (2023).

11. Wang, Y. & Wang, J. BioWorkflow: Retrieving comprehensive bioinformatics workflows from publications. Brief Bioinform 26, bbaf571 (2025).

12. Clarke, D. J. B. et al. Playbook workflow builder: Interactive construction of bioinformatics workflows. PLOS Computational Biology 21, e1012901 (2025).

13. Lee, Y. et al. Meta-Harness: End-to-End Optimization of Model Harnesses. Preprint at 10.48550/arXiv.2603.28052 (2026).

14. Lin, J. et al. Agentic harness engineering: observability-driven automatic evolution of coding-agent harnesses. Preprint at 10.48550/arXiv.2604.25850 (2026).

15. Pan, L., Zou, L., Guo, S., Ni, J. & Zheng, H. Natural-Language Agent Harnesses. Preprint at 10.48550/arXiv.2603.25723 (2026).

16. Bui, N. D. Q. Building Effective AI Coding Agents for the Terminal: Scaffolding, Harness, Context Engineering, and Lessons Learned. Preprint at 10.48550/arXiv.2603.05344 (2026).

17. Twist, L. et al. A Study of LLMs’ Preferences for Libraries and Programming Languages. Preprint at 10.48550/arXiv.2503.17181 (2025).

18. Merkel, D. Docker: lightweight Linux containers for consistent development and deployment. Linux J. 2014, 2 (2014).

19. Soiland-Reyes, S. et al. Packaging research artefacts with RO-Crate. Data Science 5, 97–138 (2022).

20. Stonebraker, M. & Rowe, L. A. The design of POSTGRES. In Proceedings of the 1986 ACM SIGMOD International Conference on Management of Data 340–355 (ACM, New York, 1986).

21. Mekonnen, A. M., Seong, K., Kim, H. & Park, J. Variant-aware Cas-OFFinder: web-based in silico variant-aware potential off-target site identification for genome editing applications. Nucleic Acids Res 53, W118– W124 (2025).

22. Patel, H. et al. nf-core/rnaseq: nf-core/rnaseq v3.22.2 - Perfect Palladium Penguin. Zenodo 10.5281/zenodo.17909656 (2025).

23. Lee, J. et al. Epithelial WNT secretion drives niche escape of developing gastric cancer. Mol Cancer 25, 1 (2025).

24. Wu, D., Gordon, C. K. L., Shin, J. H., Eisenstein, M. & Soh, H. T. Directed Evolution of Aptamer Discovery Technologies. Acc. Chem. Res. 55, 685–695 (2022).

25. Thorvaldsdóttir, H., Robinson, J. T. & Mesirov, J. P. Integrative Genomics Viewer (IGV): high-performance genomics data visualization and exploration. Brief. Bioinform. 14, 178–192 (2012).

26. Suetake, H., Fukusato, T., Igarashi, T. & Ohta, T. Workflow sharing with automated metadata validation and test execution to improve the reusability of published workflows. Gigascience 12, giad006 (2023).

27. Yao, S. et al. ReAct: Synergizing Reasoning and Acting in Language Models. Preprint at 10.48550/arXiv.2210.03629 (2023).

28. Anthropic PBC. Channels reference. Claude Code Docs. https://code.claude.com/docs/en/channels-reference (2026).

29. Robinson, J. T., Thorvaldsdottir, H., Turner, D. & Mesirov, J. P. igv.js: an embeddable JavaScript implementation of the Integrative Genomics Viewer (IGV). Bioinformatics 39, btac830 (2023).

30. Okamura, D. M. et al. Spiny mice activate unique transcriptional programs after severe kidney injury regenerating organ function without fibrosis. iScience 24, 103269 (2021).

31. Tuerk, C. & Gold, L. Systematic Evolution of Ligands by Exponential Enrichment: RNA Ligands to Bacteriophage T4 DNA Polymerase. Science 249, 505–510 (1990).

32. Ellington, A. D. & Szostak, J. W. In vitro selection of RNA molecules that bind specific ligands. Nature 346, 818–822 (1990).

33. Park, J. Towards a transparent and reproducible AI-assisted research paper writing. Genomics Inform 23, 26 (2025).

